# Effect of 2,4-Dinitrophenol Against Multidrug-Resistant Bacteria

**DOI:** 10.1101/2024.05.13.593958

**Authors:** Pérez Carrillo Víctor Hugo, Nuñez Vázquez Ángel Alfredo, Ortega Muñoz Raquel, Jiménez-Castellanos Juan Carlos, Montiel Aguirre Jesús Fernando

## Abstract

Multidrug-resistant bacteria have become very important in recent decades due to the health problems they cause. Different mechanisms of resistance in bacteria have been identified among which are efflux pumps. It is believed that chemical compounds such as 2,4-dinitrophenol (2,4-DNP) can inhibit efflux pumps which normally participate in detoxifying cells. In the present work antibiotic multiresistant cells were isolated. Resistance of these bacteria to different antibiotics was tested by the Kirby-Bauer method and ethidium bromide tolerance. It was assumed that they showed overexpression of efflux pumps. The effect of 2,4-DNP was tested and the antibiotic resistance and growth of the cells were measured. Although a strong inhibition effect by 2,4-DNP was observed, an unexpected variation in MICs among the samples suggests that 2,4DNP has a more complex effect on bacteria than originally assumed.

## Introduction

Tackling multidrug-resistant bacteria (MDR) has become one of the greatest challenges of the 21^st^ century and it is currently recognised by CDC (USA) and WHO as a severe threat to both animal and human health^1^. Ever since the discovery of penicillin in 1928 by Alexander Fleming, antibiotics have continuously been used to mitigate bacterial infections^2^.Over seven decades, new antibiotics have been developed and discovered not only for their use in humans but also in livestock^3^. Nonetheless, the overuse of antibiotics as growth promoters in animals, combined with poor clinical stewardship in humans, has exerted a strong selected pressure leading to the arise of MDR-bacteria across the world^1^. The mechanisms by which bacteria overcome the effect of antibiotics are diverse but one that stands up is the use of efflux pumps^4^. Currently, development of novel antibiotics against bacterial infections usually fails due to the inability to achieve effective antibiotic concentrations inside the bacterial cell, which is highly dependent on the permeability of the outer membrane in combination with the activity of efflux pumps^4^.

Efflux pumps are present in all bacteria and while at physiological level are actively extruding compounds, only their constitutive overproduction via loss-of-function mutations in transcriptional regulators or stress response has been linked to clinically relevant antibiotic resistance. Notably, these systems are capable to extrude, from within the cell, a myriad of chemically unrelated compounds, including fatty acids, antiseptics, detergents, virulence factors, toxins and most importantly, antibiotics^4^. Despite the extensive literature available, many questions remain open about their biology; for example, we still have not yet fully understood how they can simultaneously and almost immediately extrude a multitude of chemically unrelated substrates (e.g., antibiotics) with such an outstanding level of efficiency, speed and polyspecificity. In a great effort to combat MDR mediated by efflux pumps, different families of inhibitors have been developed such as the widely known PAβN, MBX series, NMP, CCCP, D-13 and 2,4-DNP^5,6^. The latter, is an inhibitor which acts by disrupting the proton gradient of bacterial membranes in a similar manner as CCCP inhibitor^5,7^. Although the efflux pump inhibition activity of 2,4-DNP has primarily been investigated in *Mycobacterium tuberculosis*, its phenotypic role in other clinically relevant bacteria remains not fully understood. In this work, we sought to investigate the effects of 2,4-DNP against 6 multidrug-resistant bacteria isolated from asymptomatic individuals.

## Materials and methods

### Sampling collection

We collected samples by i) swabbing the sclera and eyelids from young asymptomatic volunteers and ii) stool from also young asymptomatic volunteers. Samples were grown in Broth heart infusion media (BHI) overnight at 37 °C.

### Bacteria isolation and purification

Grown collected samples were streaked onto either MSA agar (to separate Gram-positive bacteria) or MacConkey agar (to separate Gram-negative bacteria). Plates were incubated overnight at 37 °C and fruitful isolated colonies were picked to further being biochemically characterized.

### Biochemical test for bacterial identification

We characterized bacterial population according to the classical standards for bacteria identification described by Bergey’s Manual of Determinative Bacteriology^8^

### Ethidium bromide accumulation studies

Accumulation of ethidium bromide (EtBr) was carried at different concentrations ( 0 to 100 mg/L). Bacterial load was normalized to an initial OD600=0.001. Readouts were collected at different time points by determining the presence or absence of bacterial growth (tolerance) to EtBr which is an established substrate for efflux pumps.

### Minimum inhibitory concentration (MIC)

Antibiotic disc susceptibility testing was performed according to CLSI methodology^9^. Strains were grown in the presence or absence of 2,4-DNP at a final concentration of 0.003M or 0.0005M for either Gram-negative or Gram-positive bacteria, respectively.

## Results and discussion

### Bacterial identification for collected isolates

After initial isolation on specific media, we observed that from all the samples collected, we isolated a total of 3 Gram-positive bacteria as well as 3 Gram-negative, namely **T10, WT5, A5** (for G+) while G(-) were labeled as **Vs, Ba, and Es**. Followed biochemical tests, we determined that our 3 G(+) strains were *Staphylococcus aureus* while all the Gram (-) were identified as *Escherichia coli* (Table 1). Having observed that the majority of bacteria belong to the genus *E*.*coli*, we predicted that the possibility of resistance via efflux pumps, in particular AcrAB-TolC, was almost certain according with multiple studies^10^.

**Table 1.** Biochemical characterisation of collected samples.

| Biochemical test | Samples |  |  |  |  |  |
| --- | --- | --- | --- | --- | --- | --- |
|  | **T10** | **W5** | **A5** | **Vs** | **Ba** | **Es** |
| Catalase | + | + | + | ND | ND | ND |
| Glucose | ND | ND | ND | + | + | + |
| Lactose | ND | ND | ND | + | + | + |
| Sulfuric acid | ND | ND | ND | - | - | - |
| Indol | ND | ND | ND | + | + | + |
| Motility | ND | ND | ND | + | + | + |
| Citrate | ND | ND | ND | - | - | - |
| Urea | ND | ND | ND | - | - | - |
| Manitol | ND | ND | ND | + | + | + |
| **Identified bacteria** | *S. aureus* | *S. aureus* | *S. aureus* | *E.coli* | *E.coli* | *E.coli* |

ND= Not determined

### EtBr assay shows tolerance via efflux pumps

We challenged the isolated strains to different concentrations of EtBr, a known and common substrate for efflux pumps, in a time dependent manner. Our results showed that the majority of the samples were resistant to EtBr. In particular, we observed that Gram (+) bacteria were less resistant to EtBR compared to Gram (-) (Fig 1). We surmised that the latter output is due to efflux pumps in Gram (-) bacteria, in particular those belonging to the RND-superfamily, which are more prompt to be over-expressed followed external pressure such as antibiotics.

**Figure 1.**
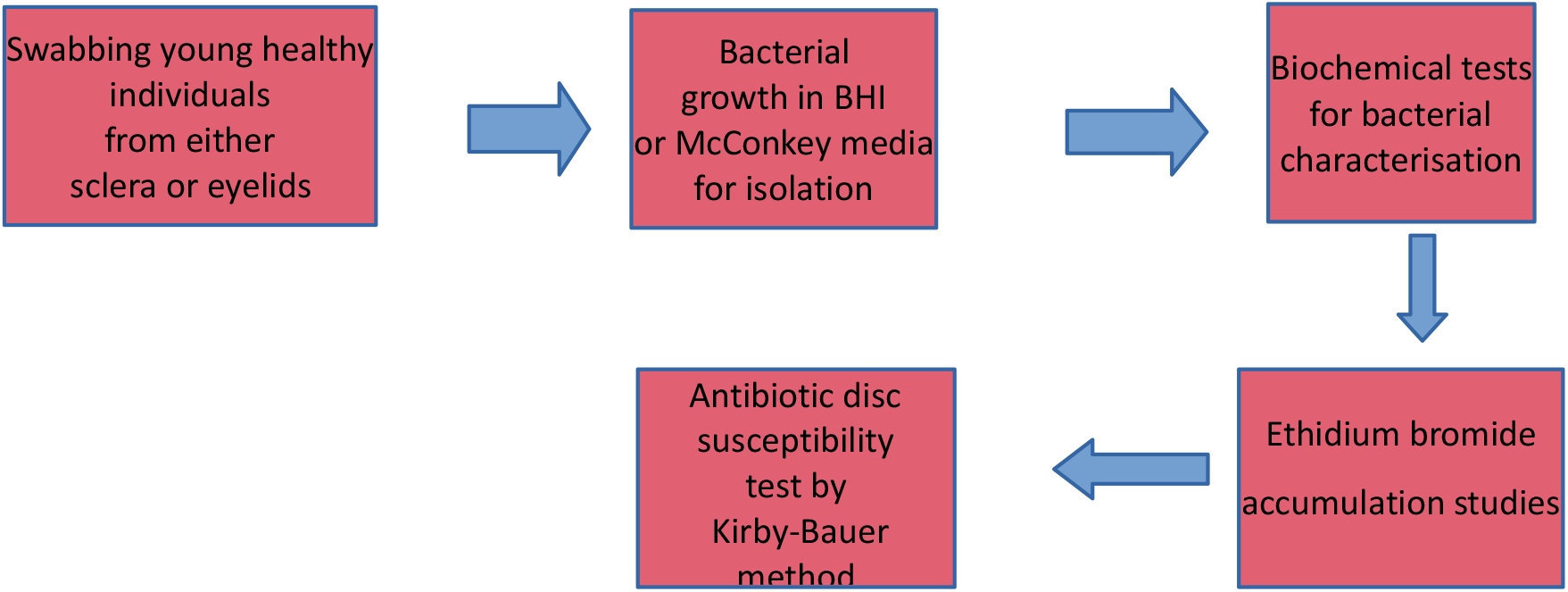
Flow chart of experiments carries in this study.

**Figure 1.**
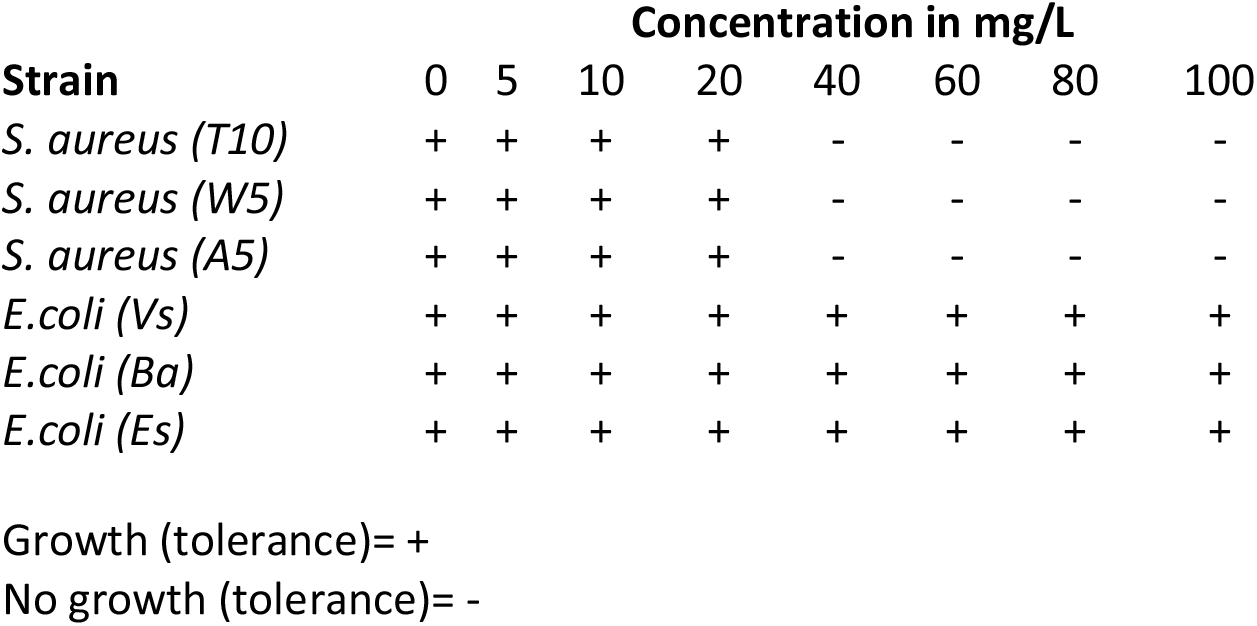
Ethidium bromide accumulation assay.

Reported values are from at least three biological repeats.

### Inhibitory effect of 2,4-DNP

We deployed solid disc MIC testing to determine the general impact of 2,4-DNP inhibition against the MDR isolates. We found that addition of the inhibitor had an approximately 25% inhibition among the antibiotics tested (Fig 2a, 2b and 2c). However, we also noticed that although some potency was achieved, a few strains (e.g., A5) showed resistance to the inhibitor 2,4-DNP (Fig 2b). It has been shown that mutations in global transcriptional regulator such as *marA* in *E. coli* or *norA* in *S. aureus* can lead to an overproduction of efflux pumps^11^. What is more, these mutations are frequently found outside healthcare settings and are easily selected in the presence of different drugs, including antibiotics. Thus, we propose that in our library, where the resistance phenotype was associated to efflux pumps (based on EtBr assays), the resistance to 2,4-DNP inhibitor (or vice versa, its lack of potency) is due to an over-expression of efflux pump via loss-of-function in key regulatory genes^12^. Alternatively, we also propose that addition of a compound that hampers the flow-through of protons, can have an overall deleterious effect not only in the membrane but also in the general bacterial physiology. Nonetheless, our findings are in accordance with the broader literature, where the direct or indirect inhibition of efflux pumps leads to a more dynamic, complex and very often confounding genotypes^12^. Together our results are helping to pave the way to test new avenues in order to continue the development of more potent efflux pump inhibitors to tackle MDR bacteria.

**Figure 2.**
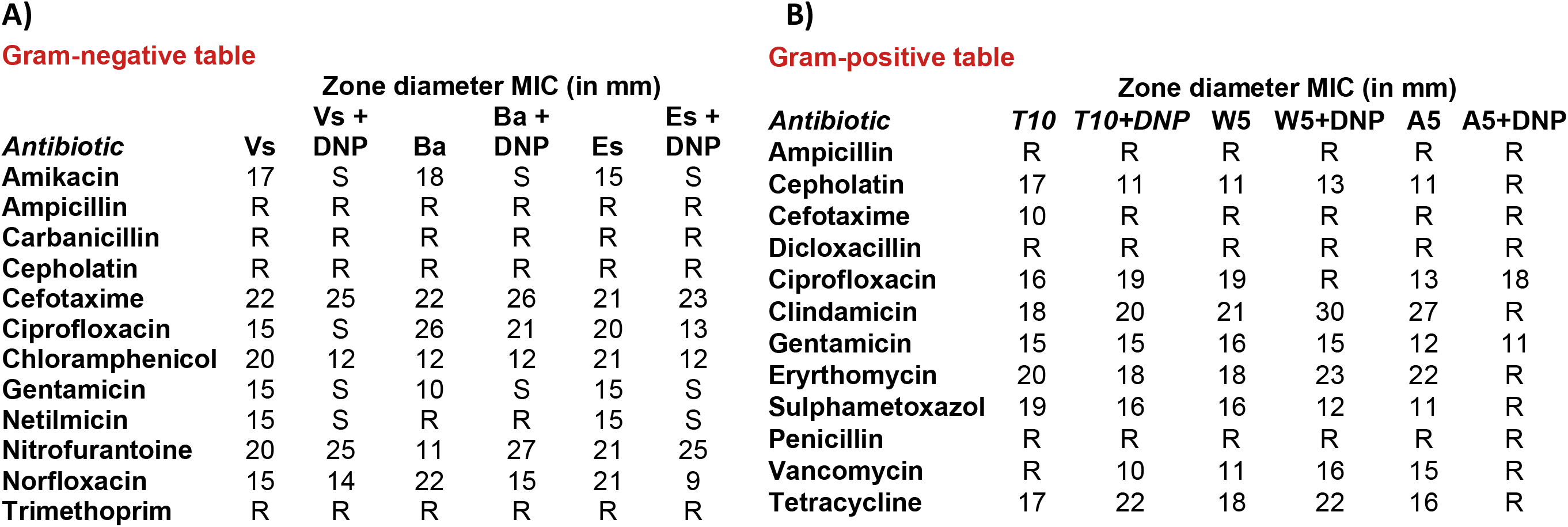
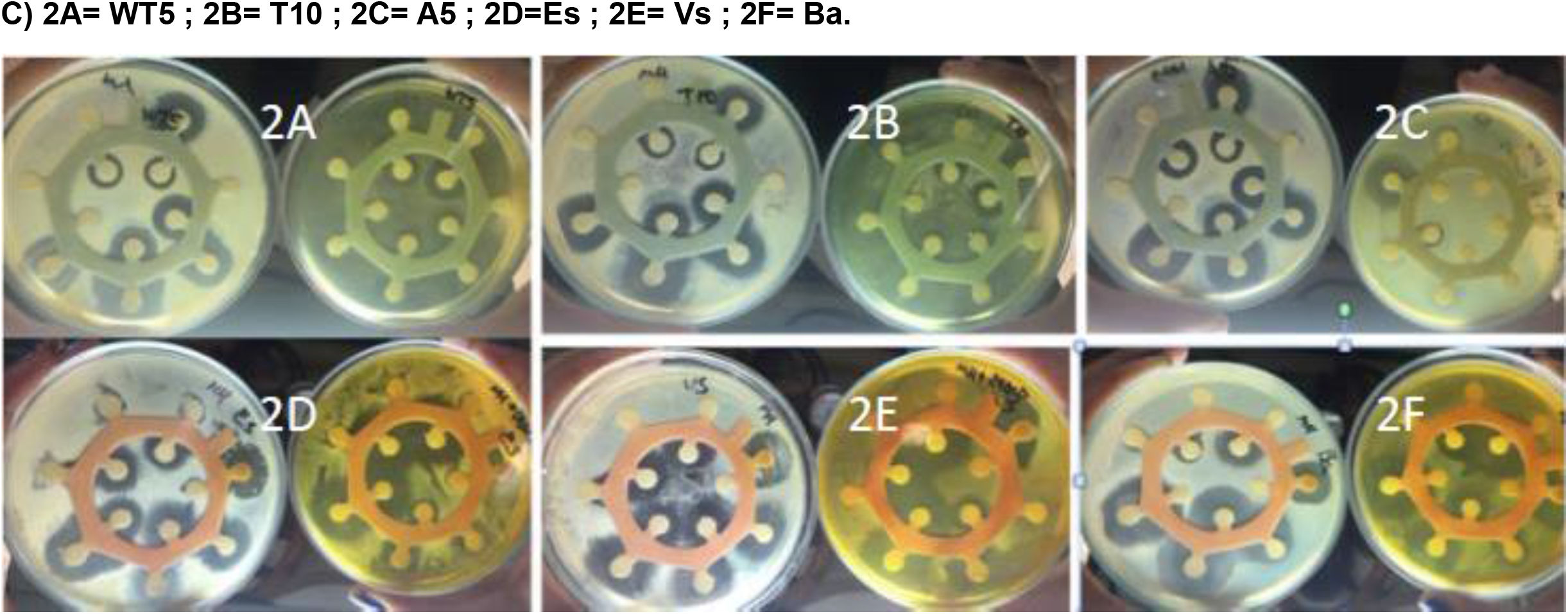
Antibiotic susceptibility disc testing (MIC) on collected MDR bacteria

## Conclusion

We isolated six multidrug-resistant strains from young asymptomatic individuals, a phenomenon that is not frequently reported. We observed that the responsibles for such phenotype belonged to either *S. aureus* and, to a greater extent, *E. coli*. Ethidium bromide tolerance assays showed that resistance is much likely driven by efflux pumps. Upon addition of 2,4-DNP inhibitor, which is a known efflux pump inhibitor, we observed an overall great antibiotic activity recovery further supporting that resistance phenotype within our isolates was due to efflux pumps activity. We observed that some strains, in particular A5, showed resistance to 2,4-DNP strongly suggesting that activity of 2,4-DNP either allows for more complex cascades for homeostatic adjustments within cell that ends helping for bacterial antibiotic resistance. The role of 2,4-DNP needs further molecular studies to gain insights about its mechanism of action and how to use this information to complement our knowledge behind the biology of efflux pump systems with the ultimate aim of tackling MDR bacteria.

## Declaration of Competing Interest

The authors declare no competing interests

## Notes

### Competing Interest Statement

The authors have declared no competing interest.

